# Multimodal Sensorimotor System in Unicellular Zoospores of a Fungus

**DOI:** 10.1101/165027

**Authors:** Andrew J.M. Swafford, Todd H. Oakley

## Abstract

Complex sensory suites often underlie critical behaviors, including avoiding predators or locating prey, mates, and shelter. Multisensory systems that control motor behavior even appear in unicellular eukaryotes, such as *Chlamydomonas*, which are important laboratory models for sensory biology. However, we know of no unicellular opisthokont models that control motor behavior using a multimodal sensory suite. Therefore, existing single-celled models for multimodal sensorimotor integration are very distantly related to animals. Here, we describe a multisensory system that controls the motor function of unicellular, fungal zoospores. We find zoospores of *Allomyces arbusculus* exhibit both phototaxis and chemotaxis. While swimming, they move towards light and settle on cellulose membranes exuding combinations of amino acids. Furthermore, we report that closely related *Allomyces* species do not share this multisensory system. Instead, each possesses only one of the two modalities present in *A. arbusculus*. This diversity of sensory suites within *Allomyces* provides a rare example of a comparative framework that can be used to examine the evolution of sensory suites. The tractability of *Allomyces* and related fungi as laboratory organisms will allow detailed mechanistic investigations into how sensory systems may have functioned in early opisthokonts before multicellularity allowed for the evolution of specialized cell types.

**Summary Statement:** Zoospores’ ability to detect light or chemical gradients varies within *Allomyces.* Here, we report a multimodal sensory system controlling behavior in a fungus, and previously unknown variation in zoospore sensory suites.

## Introduction

All organisms rely on sensory systems to gather information about their surroundings from external stimuli. The integration of individual sensory modalities into multisensory systems, or sensory suites, greatly increases the amount of information an organism can use to form responses and behaviors. Although multimodal sensory systems are common in multicellular, motile organisms, there are significantly fewer multisensory systems known from unicellular eukaryotes (Govorunova and Sineshchekov, 2005). The relative rarity of these systems in unicellular, laboratory-tractable organisms has resulted in a significant taxonomic gap between current model systems and animals.

To address this deficit, we focused on fungi characterized in part by motile, zoosporic life stages. Zoosporic fungi collectively form a clade within the ‘early-diverging lineages’ of fungi, outside the better known Ascomycota and Basidiomycota. They are largely found in freshwater ecosystems with a global distribution (James et al., 2014). Zoosporic fungi are typically characterized as saprobes, such as *Allomyces*, though parasitic life strategies on both plant and animal hosts do exist (Longcore et al., 1999). Similar to all fungi, colonies of *Allomyces* use mycelia to absorb nutrients and ultimately grow reproductive structures. Unlike most fungi, *Allomyces* produce zoosporangia, terminations of mycelial branches that make, store, and ultimately release a multitude of single-celled, flagellate zoospores (Olson, 1984). When the appropriate environmental cues are present, zoospores are produced *en masse*, eventually bursting from zoosporangia (James et al., 2014). Once in the water column, the zoospores rely on a single, posterior flagellum to propel themselves away from the parent colony and towards suitable substrates or hosts (Olson, 1984).

During dispersal of the zoosporic life stage, interpretation of environmental cues is critical for the survival and success of the future colony. Zoospores have a finite amount of endogenous energy reserves, and no zoospore is known to metabolize energy from external sources (Suberkropp and Cantino, 1973). This energetic constraint places significant pressure on the zoospore to efficiently locate a favorable environment for settlement and growth. The evolution and maintenance of a sensory system within the unicellular zoospore allows it to evaluate external conditions, move towards suitable habitats, and avoid hazards (James et al., 2014). Previous studies across zoosporic fungi have led to the discovery of a number of sensory modalities that guide zoospore dispersal including chemotaxis, phototaxis, and electrotaxis (Machlis, 1969; Morris et al., 1992; Robertson, 1972). However, these studies have neither tested a single species for multiple sensory modalities, nor posited the possibility that these single senses may only be a portion of a more complex sensorimotor system guiding zoospores.

In the fungus *Allomyces,* zoospores use chemotaxis or phototaxis to guide dispersal and settlement (Pommerville and Olson, 1987; Robertson, 1972). Chemotaxis towards the source of amino acid gradients allows zoospores to congregate at the site of an injury or on decaying material in the water column (Machlis, 1969). *Allomyces macrogynus* zoospores possess refined chemosensation, settling on substrates at varied rates in response to different amino acids (Machlis, 1969). Alternatively, the zoospores of *Allomyces reticulatus* display positive phototaxis, potentially leading spores to swim towards the air-water interface (Robertson, 1972). Studies to date have not tested for the presence of chemotaxis in *A. reticulatus* or phototaxis in *A. macrogynus*. In animals, positive phototaxis and subsequent ‘rafting’ on floating debris work to considerably increase the dispersal range of planktonic larvae (Epifanio et al., 1989). Similarly, spores attracted to the surface may encounter floating debris, algal hosts, or currents that aid their dispersal.

While little is known about the molecular mechanisms of chemotaxis in fungal zoospores, the underpinnings of their photosensitivity are beginning to come to light. The protein responsible for light-detection in zoospores of *Blastocladiella emersonii*, a close relative of *Allomyces*, is a bacteriorhodopsin gene called *CyclOps (Beme-Cycl*) (Avelar et al., 2014). Unlike many bacteriorhodopsins that regulate ion channels, *Beme-Cycl* acts through regulating intracellular cGMP (Avelar et al., 2015). A *CyclOps* gene is present in the genome of *A. macrogynus (Amag-Cycl*), a species that has been anecdotally described as having phototactic zoospores (Olson, 1984). A recent study by Gao et. al., 2015, however, contradicts claims of phototaxis in *A. macrogynus* by revealing the proteins encoded by *Amag-Cycl* are orders of magnitude less sensitive to dark/light transitions than *Beme-Cycl* proteins (Gao et al., 2015). This raises questions about the sensitivity of *CyclOps* proteins needed for phototaxis and mutations potentially responsible for shifts in photosensitivity.

The uncertainty surrounding the sensory suites of *Allomyces* zoospores demands experimental evidence to clarify number and types of modalities used during dispersal and settlement. Addressing the current deficits in our understanding of fungal sensory systems is also motivated by the potential to discover a system that will further our knowledge of multisensory evolution and function in early opisthokonts. Here, we investigate the responses to chemical and light gradients in three species of *Allomyces*; revealing previously unknown variation in fungal sensory systems, and discovering a novel multisensory system in a zoosporic fungus.

## Materials and Methods

### Culture conditions

We used *Allomyces arbusculus* str. ATCC 10983, *Allomyces reticulatus* str. California 70 from ATCC (cat. No. 42465), and *A. macrogynus* from the Roberson lab (U of AZ). We kept cultures of *A. macrogynus* and *A. arbusculus* in both solid and liquid media. For solid media, we used (Machlis, 1953) Emerson YSS (HiMedia M773) at half strength. Colonies transferred aseptically in a laminar flow hood every 4 weeks by moving a chunk of mycelia from the leading edge of the colony onto a new plate. For liquid media, we followed the protocol for Machlis’ medium B. We inoculated liquid cultures via sterile transfer of sporangia and mycelia into a 125mL Erlenmeyer flask containing 50mL of liquid media and antibiotics (Machlis, 1953) for the first generation. For all subsequent generations kept in liquid culture, we used dilute salts solution to initiate sporulation of the previous generation’s colonies. We then added 1 mL of this zoospore-dilute salts solution (referred to a sporulation product from here on) into new liquid media. Both liquid and solid cultures were grown on an orbital shaker at 140rpm and kept at room temperature (∼24C). Cultures in liquid media were kept for a maximum of 5 days, and were considered ready for sporulation after 72 hours. Because A. reticulatus did not grow well in liquid media, we cultured A. reticulatus on full strength Emerson YSS media for no more than 6 weeks. Propagation of A. reticulatus cultures in solid media was performed identically to the other species.

### Sporulation conditions

Liquid cultures of A. macrogynus and A. arbusculus were considered for sporulation after 72 hours. We visually inspected colonies under a microscope to confirm the absence of gametangia. Using a stainless steel sterile mesh, we strained the colonies out of growth media and rinsed them 5 times with dilute salts solution to remove the growth media from the colonies. We then placed the rinsed colonies and strainer in a pyrex dish with 10 mL of dilute salts solution and allowed them to sporulate for no more than 90 min. Once either sufficient spore density had been reached (5x10^5^ spores mL^-1^ for chemotaxis, 1x10^6^ spores mL^-1^ for phototaxis) or 90 minutes had elapsed, the mesh and colonies were lifted out of the dish (Machlis, 1969). Because A. reticulatus was only grown on solid media, we took a surface scraping to lift sporangia from the agar and placed it into a pyrex dish with 10 mL dilute salts solution (Saranak and Foster, 1997). If no zoospores were present, we replaced dilute salts solution every 20 minutes for the first hour. Sporulation typically occurred within 8 hours, after which the colonies were strained from the dilute salts solution.

### Phototaxis trials

We conducted phototaxis trials in a custom 1x5x3 cm (WxHxL) plexiglass chamber. We added 10 mL of sporulation product, diluted to 5x10^5^ spores mL^-1^, to the test chamber and allowed the solution 15 minutes in total darkness to dark adapt and randomize spore distribution. After dark adaption, spores were exposed to a white light (USHIO halogen bulb) through a 5 mm diameter fiber optic cable positioned 5 cm from the leading edge of the test chamber. To calibrate the amount and intensity of light, we used a JAZ Oceanoptics light sensor with Spectrasuite v2.0.162. We adjusted the intensity of the light to 1.8-1.0x10^13^ mol of photons cm^-2^ on the edge closest to the light source. Zoospores were exposed to light for 15 minutes, after which we divided the test chamber into 4 sequential sub-chambers (10x25x15mm) using sterile glass slides. This resulted in 4 sub chambers (1, 2, 3, and 4) arranged linearly so that sub chamber 1 was closest to the light source, while subchamber 4 was the farthest away (Fig. S1A). We gently agitated the liquid in each subchamber to homogenize swimming spore distribution and counted spore density in four, 10μl samples from each subchamber using a hemocyometer. A total of 10 control treatments (no light exposure) and 18 experimental treatments were conducted for each species.

### Chemotaxis trials

Chemotaxis trials followed the protocol established by Machlis (1969) (Machlis, 1969). The amino acids and combinations thereof we tested were Lysine (K), Leucine (L), Proline (P), L+K, L+P, K+P, L+K+P, and Buffer (50 mmol KH^2^PO^4^) solution (referred to as ‘treatment solutions’ from here on). All amino acid concentrations were 5x10^-4^mol. We created a chemical dispersal apparatus by drilling a hole through the lid of a 60x15mm petri dish and inserting a 5mm inner diameter glass pipette. We secured dialysis membrane (3500 MWCO) to the tip of the pipette and positioned it 3mm above the bottom of the petri dish (Fig. S1B). This creates a gradient in the petri dish of whatever solution is placed behind the dialysis membrane, allowing zoospores to navigate to the membrane, where they settle and can later be counted. The dialysis membrane was soaked and rinsed with DI water for 24 hours to remove potential contaminants and bubbles that would affect results (Carlile and Machlis, 1965).

To test for chemotaxis, we added 10 mL of sporulation product (diluted to 5x10^4^ spores mL^-1^) to the petri dish and 300 μl of treatment solution into the pipette. As a control, we used 300 μl of buffer alone. We allowed the spores to react to the gradient in total darkness for 90 minutes. At the end of the trial time, we removed the pipette and dialysis membrane from the dish and gently shook it to remove excess liquid (Machlis, 1969). We counted spores settled on the membrane under an Olympus szx7 at 400x or greater magnification.

### Molecular Methods

PCR, cDNA synthesis & sequencing: Because genomic data existed for A. macrogynus but expression data did not, we used PCR to attempt to identify if CyclOps genes are expressed in A. macrogynus zoospores. mRNA was extracted from zoospores using a NucleoSpin RNA XS kit. We synthesized cDNA using the Clontech cDNA synthesis kit. Primers were designed from putative rhodopsin/guanalyl-cyclase fusion proteins identified from the BROAD institute’s Allomyces macrogynus genome, using the CyclOps protein sequence from Blastocladiella emersonii as bait sequences (Avelar et al., 2014). Sets of primers were designed in IDT PrimerQuest. All PCRs products were visualized using a 1% agarose gel with 100bp ladder. PCR for CyclOps was only done on A. macrogynus as transcriptomes for A. reticulatus and A. arbusculus would yield expression data.

RNA was isolated from zoospores of A. reticulatus and A. arbusculus using Nucleospin xsRNA kit, and cDNA was synthesized using the NEBNext RNA First and Second Strand Synthesis modules. cDNA was sequenced using a multiplexed Illumina HiSeq lane at approximately 50x coverage.

B) Bioinformatics and Statistics: We trimmed Illumina data using Trimmomatic (Bolger et al., 2014), assembled using Trinity 2.0, and analyzed on the UCSB Osiris bioinformatics platform (Giardine et al., 2005; Oakley et al., 2014). Putative CyclOps proteins were identified using *Beme-Cycl* as a bait sequence in BLASTn searches against the *A. macrogynus* genome, NCBI bioproject 20563, and the new *A. reticulatus* and *A. arbusculus* zoospore transcriptomes. Any sequence with an e-score lower than 1e-40 was considered as a candidate. We then used the ‘get orf’ feature from Trinity to produce predicted proteins from candidate genes, selecting the longest orfs with the highest similarity to *Beme-Cycl* protein when reciprocally BLASTed using blastp. We identified the putative *CyclOps* gene in both *A. reticulatus* and *A. arbusculus* (*Aarb-Cycl*). Reads from *A. arbusculus* transcriptome were mapped back to the putative *Aarb-Cycl* using Bowtie2 (Langmead and Salzberg, 2012) and visualized using IGV viewer (Thorvaldsdóttir et al., 2013). Because the bacteriorhodopsin and guanylate-cyclase domains appeared in different ORFs on the same strand, each base was manually examined for uncertainty and low support. A guanine at site 378 was manually removed due to low coverage and low support in the reads, implying that the addition of guanine at position 378 most likely an assembly artifact. The manually edited *Aarb-Cycl* gene produced a single predicted orf with the appropriate bacteriorhodopsin-guanylyl cyclase domains (Fig. S2).

Candidate proteins were aligned using MAFFT under the L-INS-i strategy. Outgroups were selected based on a previous analysis (Porter et al., 2011) (Fig. S2). The alignment was used to create a phylogeny of candidate genes with RAxML 8 (Stamatakis, 2014) and 100 bootstrap replicates using the GTR + γ model. Trees were visualized in Evolview (He et al., 2016) and annotations were added in Adobe Illustrator.

Comparisons of zoospore phototaxis behavior were analyzed using JMP v12.0. Average spore counts per subchamber per trial were analyzed using pairwise Tukey’s HSD between control and experimental sub-chambers. Although sample size was low, each sample is an average of four replicates - making effective sample size close to 40 for each treatment. Zoospore chemotaxis was analyzed using Wilcoxon each pair due to the nonparametric distribution of results and small sample size (n=10).

## Results

### A. reticulatus relies on phototaxis

Zoospores of *A. reticulatus* showed no significant deviation from the control when exposed to any amino acid treatment (P > 0.05 for all treatments) (Fig. 1). In accordance with existing literature, *A. reticulatus* showed a significant response to a light gradient (Robertson, 1972). The number of zoospores swimming in the subchamber closest to the light source (Fig. 2) was significantly higher than when no light source was present (P = 0.0005).

**Figure 1.**
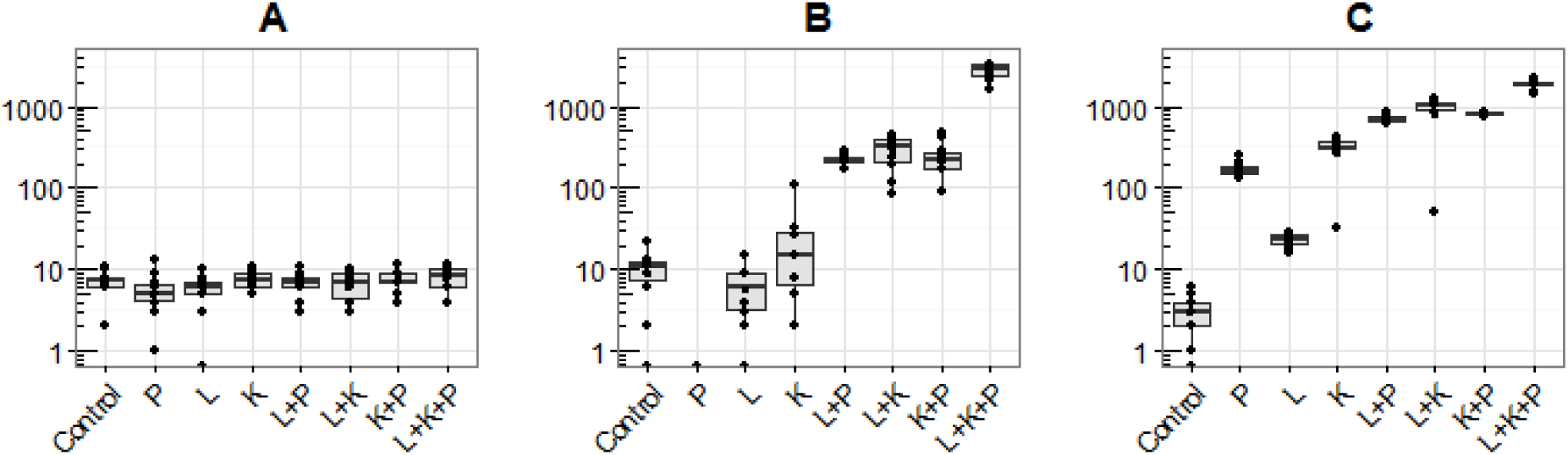
The number of zoospores settled on the dialysis membrane in response to varying amino acid treatments. (**A**) *Allomyces reticulatus,* (**B**) *A. arbusculus,* (**C**) *A. macrogynus*. Each column represents the number of zoospores settled on 2mm^2^ dialysis membrane after 90 minutes. Squares represent means. **Control**, Proline (**P**), Leucine (**L**), Lysine (**K**), and combinations thereof. N=10 for all treatments, Y axis in log scale.

**Figure 2.**
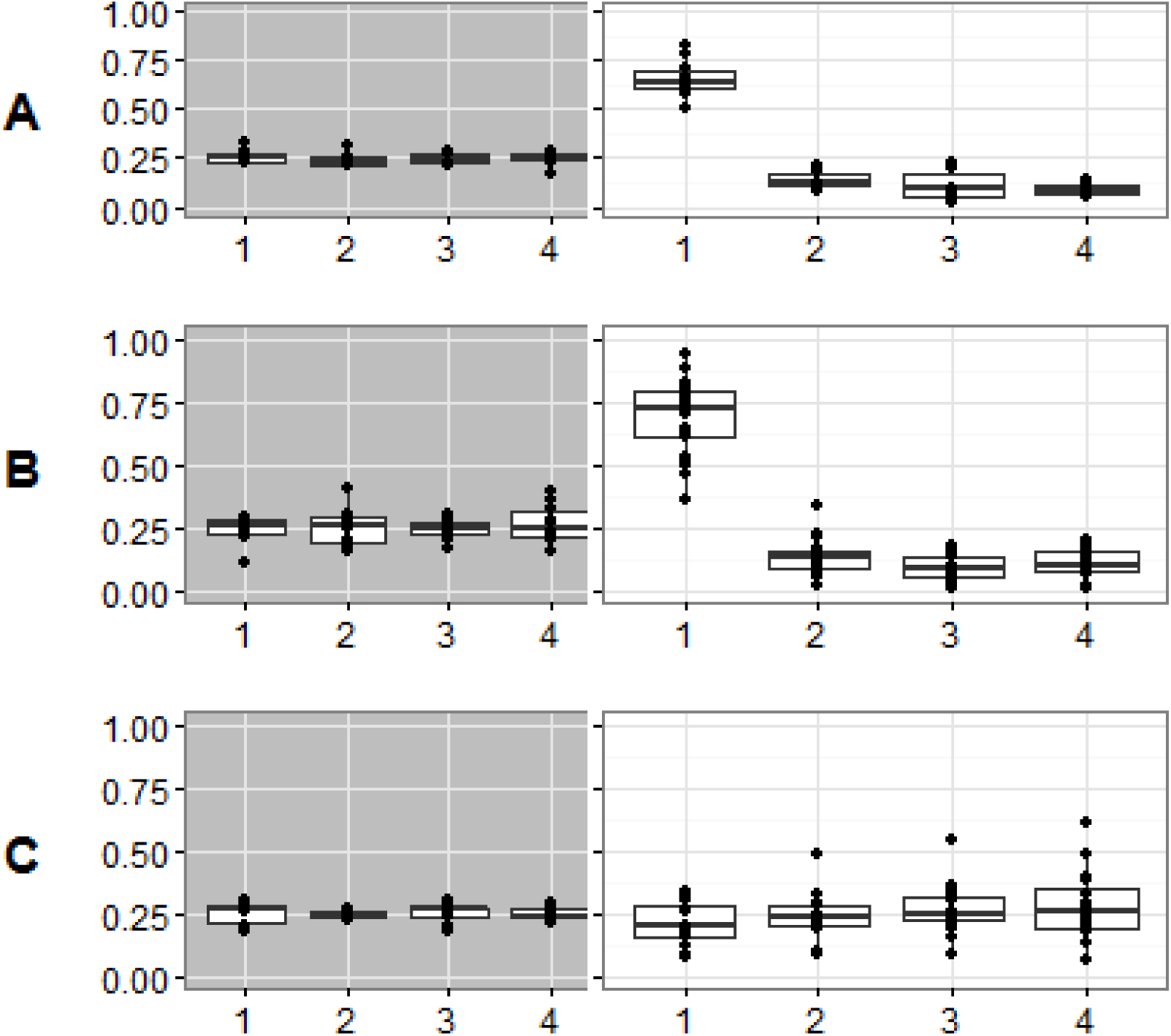
The distribution of swimming zoospores found in each subchamber in phototaxis trials, shown as percentage. (**A**) *A. reticulatus* (**B**) *A. arbusculus* or (**C**) *A. macrogynus* zoospore distribution after 30 minutes of darkness (**grey background**) or 15 minutes of darkness followed by 15 minutes exposure to directional light (**white background**). The directional light source was positioned so light intensity was strongest in sub-chamber **1** and lowest in sub-chamber **4**.

### A. macrogynus relies on chemotaxis

As seen in previous experiments, *A. macrogynus* zoospores displayed a significant response to all amino acid treatments compared to the control (Machlis, 1969). Proline (P = 0.0294), leucine (P=0.0275), lysine (P=0.0294), and any combination of two or three amino acids when compared to a control (P < 0.001) (Fig. 1). Zoospore response to increasing treatment complexity was non-linear though roughly equal for all unique combinations of equal complexity. *A. macrogynus* zoospores showed no response when exposed to a light gradient (Fig. 2). The number of zoospores in all sub-chambers were the same for both light and dark trials (P = 0.9976).

Allomyces arbusculus *uses both chemotaxis and phototaxis in a multisensory system*

As expected from existing literature, *A. arbusculus* zoospores responded similarly to the zoospores of *A. macrogynus* when exposed to amino acid treatments. The number of spores settled increased in a nonlinear fashion as the complexity of the treatment increased, though at a lower average number of settled spores when compared to *A. macrogynus*: K+P (P = 0.0014), K+L (P = 0.0014), L+P (P = 0.0008), K+L+P (P = 0.0004) (Fig. 1). As opposed to *A. macrogynus, A. arbusculus* does not respond to, or cannot detect, gradients of single amino acids (P > 0.05 for all single A.A. treatments) with the possible exception of Proline. We found that no zoospores settled when *A. arbusculus* was exposed to trials of Proline alone, potentially indicating the potential for negative chemotaxis or inhibition of settlement in response to gradients of Proline by itself. However, we can not definitively resolve this reaction with our sample size (Proline v. Control: P = 0.072; N=10). When exposed to a directional light source, *A. arbusculus* display positive phototaxis (Fig. 2). The number of zoospores in the sub-chamber closest to the light was significantly higher (P < 0.0001) in light vs. dark trials and was comparable to the response of *A. reticulatus* zoospores.

### CyclOps is present in all species, may not be expressed in A. macrogynus zoospores

Phylogenetic analysis of putative *CyclOps* genes reveals *CyclOps* presence in all three samples (*A. arbusculus & A. reticulatus*: transcriptome data. *A. macrogynus*: previously available genome data). The single copy of *CyclOps* recovered from the *A. arbusculus* transcriptome revealed a possible truncation of the Guanylyl cyclase domain (Fig. 3). Both *Aarb-Cycl* and *Amag-Cycl* share a mutation at the putative functional residue F313 to I313. Despite the success of positive controls indicating successful PCR amplification, no primers successfully amplified *Amag-Cycl* from *A. macrogynus* zoospores.

**Figure 3.**
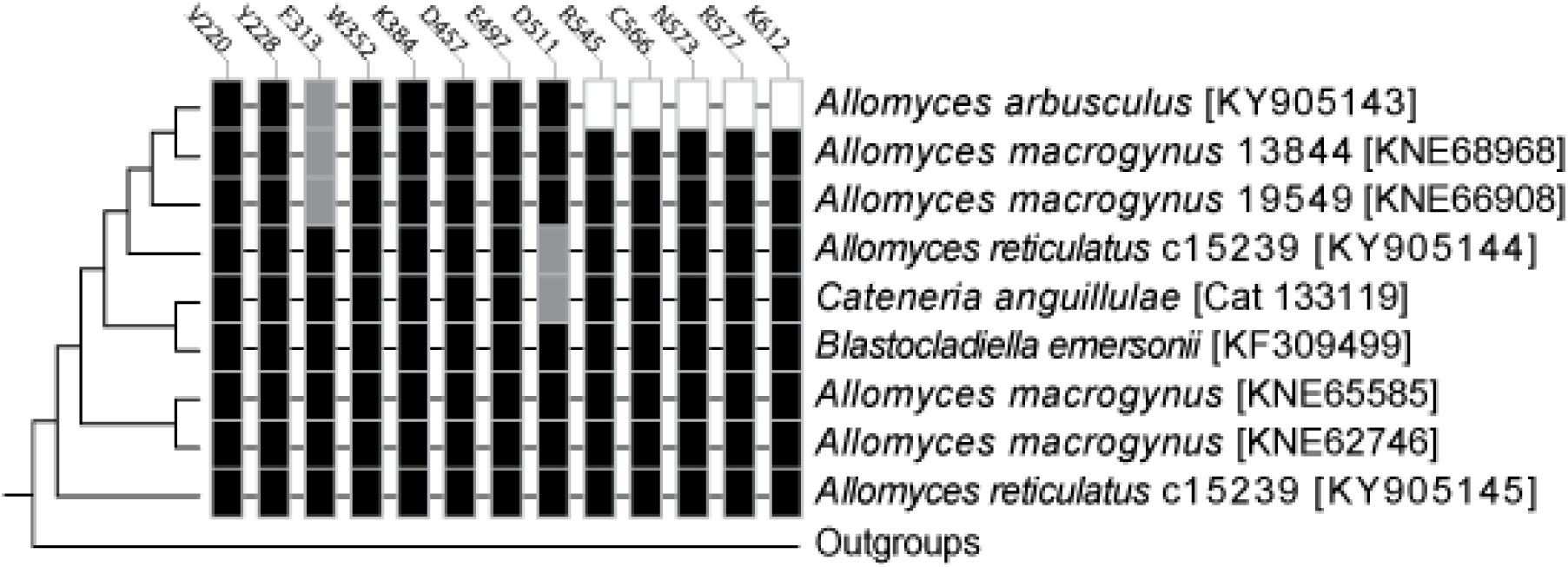
Cladogram showing the relationships and conserved amino acid residues in CyclOps proteins. Boxes indicate amino acid residues critical for binding (Avelar et al., 2014), with residue numbers based on position in unaligned Beme-Cycl (AIC07007.1). Black boxes indicate an amino acid matching at that residue when compared to Beme-Cycl, grey boxes indicate a mutation, and white boxes indicate a gap.

## Discussion

Understanding how sensory modalities evolve and integrate with other behavioral circuits remains an open question in neurobiology and evolutionary biology. The *Allomyces* genus, with new variation in sensory suites discovered in this study, will aid in answering these questions. Previous studies had shown *Allomyces* spores use either chemotaxis or phototaxis to guide dispersal. Here, we reveal that the sensorimotor system in *A. arbusculus* is multimodal-able to process both chemical and light cues. Additionally, our results reveal the previously unknown complexity and variation of sensorimotor systems in *Allomyces*.

### Variation in sensory modalities across Allomyces

In this report, we discover unknown variation in the distribution of sensory modalities across the genus. This variation manifests in two ways: the types of sensory modalities used by each species of *Allomyces* and the responses of *A. arbusculus* and *A. macrogynus* zoospores to the same amino acids. The lack of phototaxis, coupled with the inability to amplify *CyclOps* from zoospore mRNA in *A. macrogynus* is quite interesting due to conflicting reports between previous studies. These studies conflictingly report that *A. macrogynus* zoospores are either phototactic (through anecdotal evidence) (Olson, 1984) or that the photosensitive protein, Amag-Cycl, does not respond effectively to light (Gao et al., 2015). Our findings support the hypothesis that *CyclOps* in *A. macrogynus* no longer effectively differentiates between light and dark, and suggest that *CyclOps* expression has been lost in zoospores. With the loss of phototaxis, the distribution and settlement of *A. macrogynus* zoospores likely deviated greatly from both *A. reticulatus* and *A. arbusculus* in natural settings.

The variation in *Allomyces* sensory suites discovered in this study coupled with the convergent function of *CyclOps* and animal opsins make it critical to our understanding of the evolutionary history of light sensing. Animal photoreception, mediated through Type II opsins, operates through intracellular regulation of cyclic nucleotides. Non-animal photoreception, mediated through Type I opsins, operates through channel and sensory rhodopsins (SRII). SRIIs, like type II opsins, induce a signal transduction cascade. However, unlike animal opsins, SRIIs do not regulate cyclic nucleotide concentrations. Instead, SRIIs indirectly regulate CheY, a protein that controls flagellar motion (Klare et al., 2004). Type I and II opsins are considered a spectacular example of convergent evolution (Larusso et al., 2008). *CyclOps* represents a third case of convergence, where protein function may modulate intracellular cGMP levels, yet the sequence similarity of *CyclOps* imply its origin as a diversification of type I opsins (Avelar et al., 2015). Studying the evolution of *CyclOps* sequence function through the variation in *Allomyces* will yield further insights into the evolution of novel photosensory mechanisms.

The lack of chemotaxis in *A. reticulatus* highlights the variation in *Allomyces* sensory system evolution. While previous studies uncovered differences in the combinations of amino acids *A. arbusculus* and *A. macrogynus* zoospores prefer (Machlis, 1969), no one tested the possibility of a complete lack of chemotaxis in a close relative. The discovery of *A. reticulatus’s* lack of chemotaxis reveals a turnover in sensory modalities responsible for controlling a vital behavior across the genus. Future studies will use the variation in both chemotaxis and phototaxis across *Allomyces* as a platform to understand behavioral integration, multisensory evolution, and sensory remodeling in an organism closely resembling ancestral opisthokonts.

### The Multisensory System of Allomyces arbsuculus

*Allomyces arbusculus* zoospores present an easily culturable, laboratory tractable system for investigating multimodal sensation in unicellular systems and its underlying mechanisms. Previous studies have independently confirmed that zoospores use a diversity of senses, but most zoospores have only been tested for a single sense (Avelar et al., 2014; Machlis, 1969; Morris et al., 1992; Robertson, 1972). Our findings represent the first study of zoospore multisensory capabilities, and a concrete example of multimodal sensorimotor control in a unicellular opisthokont (Fig. 1-2). Though choanoflagellates fall within Opisthokonta (Cavalier-Smith et al., 2014) and potentially exhibit both areo-and pH-taxis, it remains unknown if the sensory modalities in colonies (areotaxis) are also used to direct the dispersal stage (pH-taxis) (Kirkegaard et al., 2016; Miño et al., 2017).

Zoospores, as unicellular, flagellated cells, might closely represent the ancestral opisthokont phenotype (Cavalier-Smith et al., 2014). Specialized cell types and functions may often evolve through subfunctionalization followed by elaboration of the ancestral cell’s functions (Arendt et al., 2016). This implies that as multicellular opisthokonts evolved, the foundation for specialized sensory modalities already existed. Under the subfunctionalization hypothesis, multimodal systems in unicellular organisms, such as we report in *A. arbusculus*, must have evolved prior to subfunctionalization in ancestral, multicellular opisthokonts. Future studies of the multimodal sensorimotor system in *A. arbusculus* zoospores and the variation in modalities across *Allomyces* may uncover how multiple senses became integrated into behavioral responses in ancestral opisthokonts. Understanding these mechanisms in the context of the cellular subfunctionalization hypothesis will further our understanding of sensorimotor evolution, elaboration and individuation through cellular specialization.

### Conclusions

We present a multimodal sensorimotor system in a unicellular life stage of a fungus. The multisensory suite of *A. arbusculus* zoospores is an excellent system to study how sensory modalities integrate into existing behavioral regimes. Together with existing models of unicellular sensory mechanisms, the variation in sensory modalities in *Allomyces* and other early diverging fungi will allow us to formulate more accurate conclusions about the evolution of complex sensory suites and multisensory systems in ancestral eukaryotes. Lastly, the relatively narrow taxonomic breadth associated with the multiple transitions in sensory suites during *Allomyces* evolution will allow testing of broad questions in evolution; such as the role multimodal cell types play in the origin and evolution of specialized traits, and how emergent behaviors evolve during sensory remodeling.

## Acknowledgements

The authors would like to thank Robert Roberson for supplying *A. macrogynus* and Jason Stajich for his advice and guidance on molecular work in fungus systems. Additionally, the authors would like to thank Oren Malvey for his help in maintaining cultures.

## Author Contributions

A.S. and T.O. designed experiments. A.S. carried out experiments and analyzed data. A.S. and T.O. wrote and edited the manuscript.

## Data Availability

Alignment, newick, and behavioral data collected for this study is available for download at https://bitbucket.org/swafford/multisensory_systems_JEB_2017.

